# Characterization of segmental duplications and large inversions using Linked-Reads

**DOI:** 10.1101/394528

**Authors:** Fatih Karaoglanoglu, Camir Ricketts, Marzieh Eslami Rasekh, Ezgi Ebren, Iman Hajirasouliha, Can Alkan

**Author notes:** Joint corresponding authors.

## Abstract

Many algorithms aimed at characterizing genomic structural variation (SV) have been developed since the inception of high-throughput sequencing. However, the full spectrum of SVs in the human genome is not yet assessed. Most of the existing methods focus on discovery and genotyping of deletions, insertions, and mobile elements. Detection of balanced SVs with no gain or loss of genomic segments (e.g., inversions) is particularly a challenging task. Long read sequencing has been leveraged to find short inversions but there is still a need to develop methods to detect large genomic inversions. Furthermore, currently there are no algorithms to predict the insertion locus of large interspersed segmental duplications.

Here we propose novel algorithms to characterize large (>40Kbp) interspersed segmental duplications and (>80Kbp) inversions using Linked-Read sequencing data. Linked-Read sequencing provides long range information, where Illumina reads are tagged with barcodes that can be used to assign short reads to pools of larger (30-50 Kbp) molecules. Our methods rely on *split molecule* sequence signature that we have previously described [11]. Similar to the split read, split molecules refer to large segments of DNA that span an SV breakpoint. Therefore, when mapped to the reference genome, the mapping of these segments would be discontinuous. We redesign our earlier algorithm, VALOR, to specifically leverage Linked-Read sequencing data to discover large inversions and characterize interspersed segmental duplications. We implement our new algorithms in a new software package, called VALOR_2_.

**Availability:** VALOR_2_ is available at https://github.com/BilkentCompGen/valor.

## 1 Introduction

Alterations of DNA content and organization larger than 50 bp, commonly referred to as genomic structural variations (SVs) [2], are among the major drivers of evolution [24, 29], and diseases of genomic origin [38]. Despite decades of research they remain difficult to accurately characterize contributing to our lack of full understanding of the etiology of complex diseases, termed *missing heritability* [9].

High-throughput sequencing (HTS) technologies are widely employed to discover and genotype various classes of SVs since their inception [18, 13, 26, 34, 12, 19, 36]. However, effectiveness has been limited by either very short read lengths (e.g., Illumina), or high error rates and prohibiting cost (e.g., PacBio and Oxford Nanopore). The human genome complexity further contributes to our lack of full characterization of structural variants, especially large-scale duplications and balanced rearrangements due to the repetitive and duplicated sequence at the SV breakpoints [17]. Despite high error rates, long reads offer improvement in complex SV discovery, either used alone [10, 16], or when integrated with standard short-read sequencing data [32].

Recently Linked-Read sequencing methods such as the 10x Genomics system (10xG) was introduced as an alternative method to generate highly accurate Illumina short reads data with additional long-range information [27]. In the 10xG system, large DNA molecules (typically 10-100 Kbp) are barcoded and randomly separated into over a million partitions (here we term these partitions “pools”). Each pool contains roughly 2-30 large molecules. These pools are then sequenced at *very low coverage* (∼0.1X) using the standard Illumina platform. Shared barcodes among Illumina read pairs show them as generated from the same pool. Since each pool is diluted to contain only a very small fraction of the input DNA, the probability of barcode collision is negligible [45]. For example, assuming 20 molecules per pool and an average size of 30 Kbp per molecule, each pool on average contains only 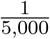 of the haploid human genome. Linked-Reads then can be used to “reconstruct” large molecules that originate from the same haplotype. Furthermore, Linked-Read sequencing makes it possible to obtain very high physical coverage with the cost of generating moderate sequence coverage data^1^.

The ability of extracting long range information from accurate and inexpensive but short read sequencing data makes Linked-Read sequencing attractive for various applications. It has been used for genome scaffolding [47], haplotype-aware assembly [27, 33, 43], metagenomics [8], single cell transcriptome profiling [35, 44] and regulatory network clustering [1], haplotype phasing [27, 33, 48], and genome structural variation discovery [11, 23, 37, 45].

Linked-Read techniques for genomic structural variation discovery include VALOR [11], Long Ranger [23] and GROC-SVs [37]. VALOR was the first algorithm that used “split molecule” signature, similar to the commonly used split read signature [46], together with traditional read pair signature [42, 26, 2] to characterize large (>500 Kbp) inversions. Split molecules are defined as large molecules that span an SV breakpoint, and therefore mapped as two disjoint intervals to the reference genome.

Long Ranger[23] is a comprehensive software package developed by 10X Genomics, for the purpose of barcode-aware read alignment and resolving full-scale human germline genome variation, while GROC-SVs is an optimized tool for somatic and complex SVs in cancer genomes. Both Long Ranger and GROC-SVs employ a novel idea to utilize discordance in expected “barcode coverage” as well as barcode similarities across distant locations for potential large-scale SV signals. In addition, GROC-SVs [37] performs local assembly on barcoded reads to detect large complex events that are between 10-100 Kbp with breakpoint resolution.

Despite the aforementioned advances in SV discovery using various technologies, detecting both balanced rearrangements (i.e., inversions and translocations) and segmental duplications (SDs) remain challenging due to mapping ambiguity. Note that it is still possible to identify increase in SD copy number using read depth signature [3, 40], however, no method yet exists to *anchor* a new SD (i.e. find their insertion locations).

Here we present **novel algorithms** to discover large (> 40Kbp) direct and inverted interspersed SDs using Linked-Read sequencing data. We redesign and extend upon VALOR and use split molecule and read pair signatures to detect SDs and estimate the insertion sites of the new SD paralogs, and further include read depth signature to filter potential false positives caused by incorrect mappings. We implemented our new algorithms as the VALOR_2_ software package. Briefly, VALOR_2_ differs from the former version of VALOR through: 1) it can characterize segmental duplications in both direct and inverted orientation, 2) incorporates read depth information to improve predictions and reduce false calls, and 3) provides full support to alignment files (i.e., BAM) generated from 10xG Linked-Read data sets.

Using simulated data sets we show that VALOR_2_ achieves high precision and recall (94% and 82%, respectively) for segmental duplications, and 98% and 76% for large inversions. We also applied VALOR_2_ to the genome of NA12878 sequenced with the 10xG platform [27] and identified 5 direct, and 9 inverted segmental duplications.

## 2 Methods

We have previously described an earlier version of VALOR_2_ that uses split molecules and read pair signature to detect inversions [11]. Here we describe novel formulations, algorithms and optimizations to characterize large (> 80Kbp) inversions and (> 40Kbp) *segmental duplications* in both direct and inverted orientation. We depict the split molecule and read pair sequence signatures for these types of large SVs in Figure 1.

**Figure 1:**
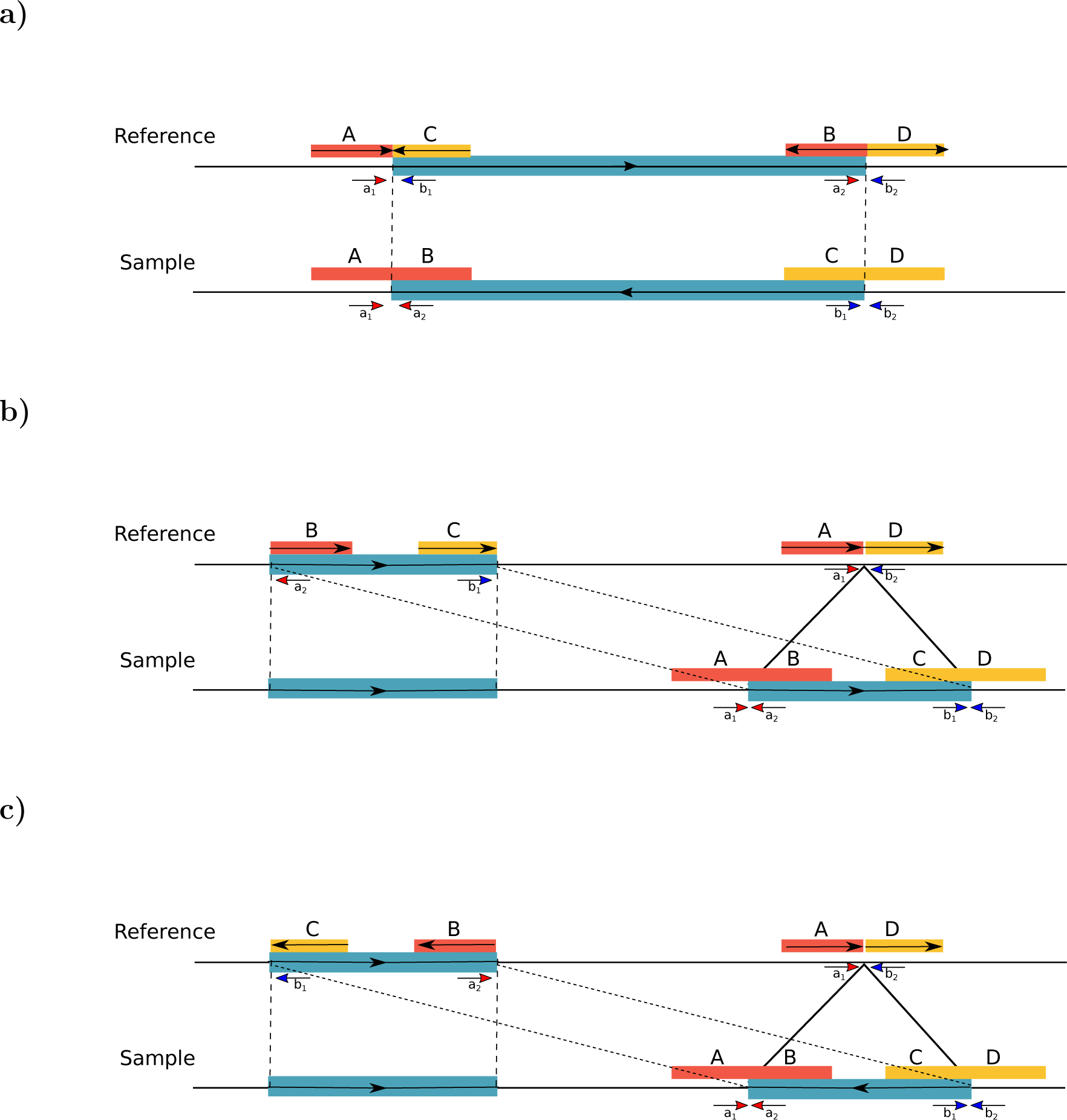
Split molecule and read pair sequence signatures used in VALOR_2_. a) Inversion, b) interspersed duplication in direct orientation, c) inverted duplication. In each case, the large molecules that span the SV breakpoints are split into two mapped regions. Note that, it is not possible to determine the mapped strand of the split molecules shown here. From the perspective of the reference genome (i.e., mapping), A,B,C,D are defined as *submolecules*, A/B and C/D pairs are *candidate splits*, and A/B-C/D quadruple is a *split molecule pair*.

### 2.1 Glossary

Here we define several terms that we use in this manuscript:

- *molecule*: a large molecule (30-50 Kbp) that was barcoded and pooled using the 10xG platform. Here we refer to the physical entity.
- *submolecule*: a molecule identified *in silico* by the VALOR_2_ algorithm by analyzing read map locations.
- *candidate split*: a pair of submolecules with the same barcode that potentially signal a SV event.
- *split molecule pair*: a pair of candidate splits with different barcodes that potentially signal the same SV event.

### 2.2 Overview of the VALOR_2_ algorithm

VALOR_2_ depends on only the alignment files (i.e., BAM) with the necessary barcode information generated with Long Ranger, BWA-MEM, or a similar read mapper. Briefly, VALOR_2_ first tries to identify the underlying large molecules separately for each barcode, which we call *submolecules*. In this step, we do not consider reads that map to satellite regions, and we discard very short submolecules. Two identified submolecules are paired together (called *candidate splits*) if the summation of their span is *≤ µ*_molecule_ + 3*σ*_molecule_ where *µ*_molecule_ is the average and *σ*_molecule_ is the standard deviation of the inferred submolecule sizes. Next, VALOR_2_ removes those candidate splits with no read pair support. It also discards those that signal a duplication event without read depth support. Additionally, any candidate splits that span assembly gaps are removed from consideration. VALOR_2_ then 1) pairs candidate splits with different barcodes that likely signal the same SV event (*split molecule pairs*), and 2) models the split molecule pairs as vertices in a graph and solves the maximal quasi clique problem [6]. In this graph, edges represent overlap (i.e., “agreement”) between two split molecule pairs. Finally, VALOR_2_ reports SVs that are supported by at least two pairs of split molecules.

Below, we present the details for each step in the VALOR_2_ algorithm.

### 2.3 Molecule Recovery

The first step of the VALOR_2_ algorithm involves identification (or, recovery) of the large molecules from mapped data. Initially, we call the intervals returned by this recovery as *submolecules*. For this purpose we use a sliding window approach to greedily group reads with the same barcode that are mapped in close proximity (Algorithm 1). Here we only consider concordantly mapped read pairs, and we take the full span of a read pair as a *fragment*. For each barcode, we scan each chromosome and merge together fragments if they are within a user-defined distance *T*, or if a new fragment is within distance *Q* from the leftmost fragment in a re-identified submolecule. We use *Q* = 80, 000 and *T* = 10, 000 by default^2^, determined by parameter sweeping. Finally, we remove very short submolecules (*<*3 Kbp by default) that correspond to less than 1/10 of expected average molecule size from consideration.

### 2.4 Clustering using SV graph

We first record all pairs of submolecules that share the same barcode and map to the same chromosome as *candidate splits*, and then compare all possible pairs of candidate splits across different barcodes to find those that signal an inversion or a duplication, termed *split molecule pairs* (see Figure 1 for the depiction of candidate splits and split molecule pairs). We limit inversion predictions and the duplication size by the largest inversion size we can find in the literature [4] (≈7 Mbp). Next, we test whether the split molecule pairs are supported by read pair signature (Figure 1). Here we require at least 3 read pairs to signal the same SV event and we remove candidate splits with insufficient support from consideration.

We construct an SV graph *G* as follows (Figure 2). We denote each remaining split molecule pair as a vertex in *G*, and we create an edge between two vertices if their corresponding split molecule pairs signal the same SV event. Finally on the resulting graph we find clusters of read pair supported split molecule pairs by approximately solving the maximal clique problem using the quasi-clique formulation [6]. Here a quasi clique is defined as an approximate clique with *V* vertices and 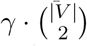 edges, where *γ* is a user-defined parameter, which we set to *γ* = 0.6 by default. Each quasi clique defines a putative SV event.

#### Algorithm 1 Molecule recovery.

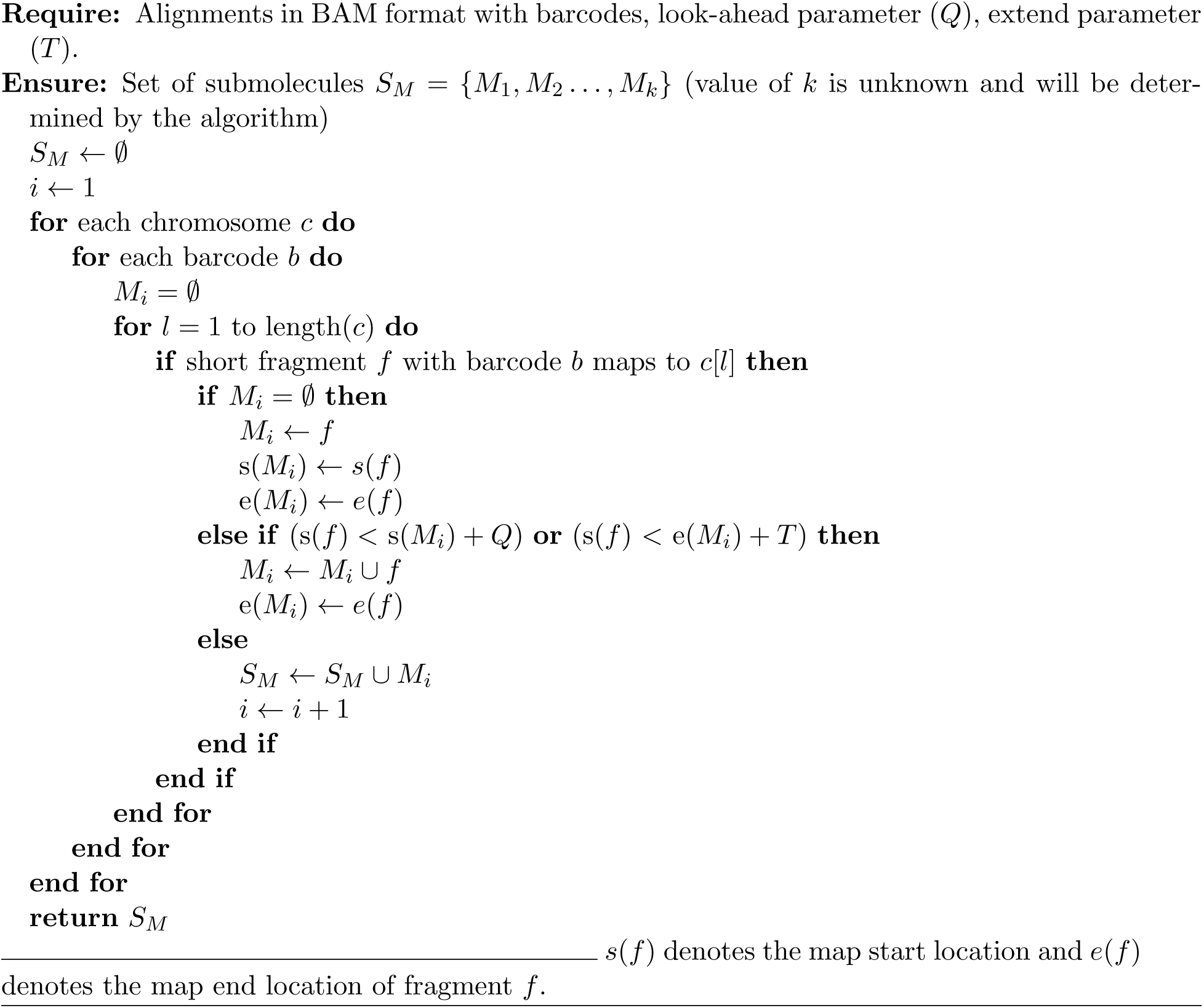

**Figure 2:**
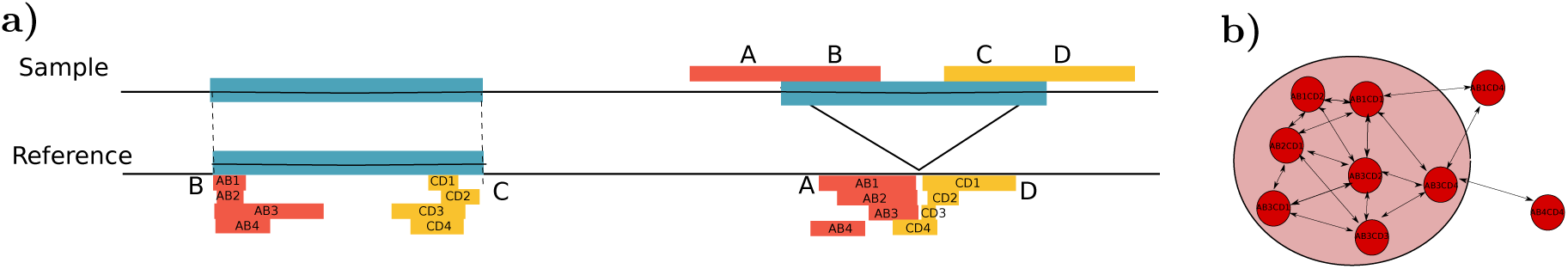
Building the SV graph from split molecule pairs. a) Four pairs of split molecules that signal a segmental duplication. b) Corresponding SV graph, where each vertex denotes a pair of submolecules that signal the duplication, and edges show “agreement” between pairs. The shaded area corresponds to the quasi-clique selected as representative of the putative SV.

We identify inversions breakpoints with two coordinates, and duplications with three coordinates. The third coordinate for a duplication event denotes the insertion breakpoint, for which we provide a confidence interval.

### 2.5 Molecule depth filtering

Although there are only a small number of molecules that share the same barcode (2-30), it is still possible that two or more different molecules originate from the same chromosome. Additionally, the molecule sizes do not follow Gaussian, Poisson, or a similar distribution (Figure 3), thus it is not possible to distinguish true split molecules from “normal” but short molecules. The read pair sequence signature is not entirely reliable either due to the mis-mapping artifacts within or around repeats and duplications. We, therefore, apply additional filtering on duplication calls based on “molecule depth”. We reason that the number of molecules that originate from segmental duplications must be higher than the genome-wide average, similar to the traditional read depth signature [5, 3]. In this step, we first calculate the average molecule depth (*µ*_depth_) and standard deviation (*σ*_depth_) in the entire genome. We then discard segmental duplication predictions with molecule depth *< µ*_depth_ + *σ*_depth_.

**Figure 3:**
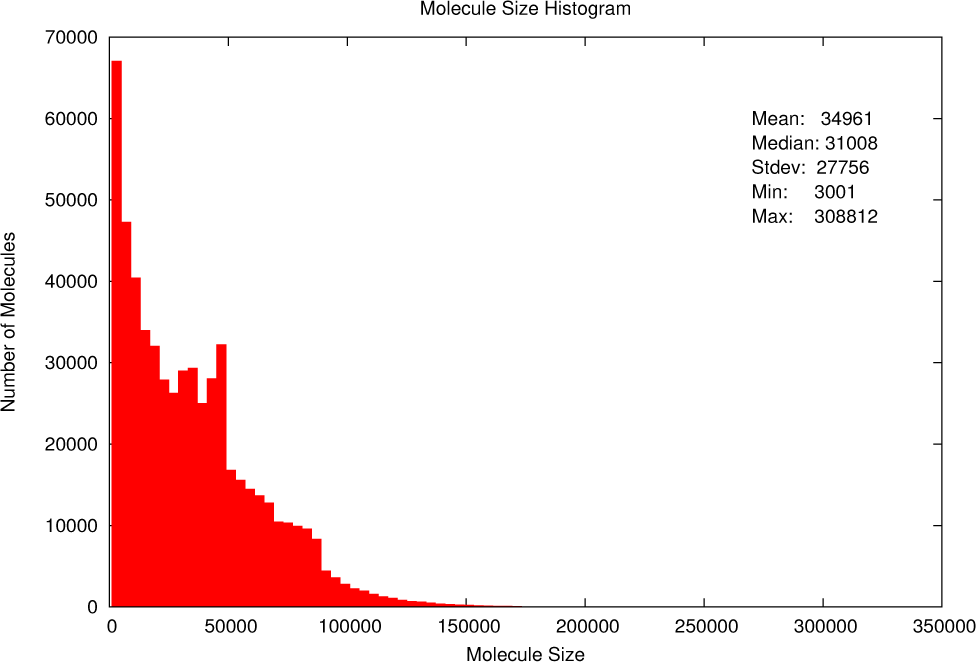
Molecule size histogram mapped to chromosome 16 as observed in the Linked-Read sequencing data generated from the genome of the NA12878 sample [27].

## 3 Results

We tested VALOR_2_ using both simulated and real data sets to compare the precision and recall rates of VALOR_2_ with one other tool that use Linked-Read sequencing (Long Ranger [23], and two tools that use only WGS data sets (DELLY [31] and LUMPY [19]). However, VALOR_2_ is the only tool that can characterize interspersed duplications, therefore we limit our comparison to only inversions, and evaluate VALOR_2_’s performance on duplications using simulations. We find that VALOR_2_ is complementary to other methods in inversion calls as VALOR_2_ aims to find larger (>80Kbp) inversions, while the other tools focus on smaller (*<*100 Kbp) SVs.

### 3.1 Simulation experiments

We used VarSim [28] to generate a simulated diploid human genome. Our simulation included variants of various lengths and types: 2.8 million SNPs, ≈195,000 indels, and ≈ 5,000 SVs (>50 bp, up to 6 Mbps). We found that VarSim only generates tandem duplications, therefore we randomly changed a subset of simulated tandem duplications to interspersed in the simulated VCF file, assigned random insertion breakpoints, and then applied changes to the reference. We then generated Illumina WGS reads at 40X depth of coverage using ART [14], and 10xG Linked-Reads at 50X coverage using LRSim [22]. The 10xG Linked-Reads simulation has extra coverage to account for the barcode sequences that are part of the read and other losses as also described in [23].

We mapped the simulated reads to the human reference genome (GRCh37) using BWA-MEM [20] for WGS, and Long Ranger for 10xG data sets. We then applied the standard BAM processing that includes sorting with SAMtools [21] and marking duplicates with Sambamba [41]. We used VALOR_2_ and Long Ranger to generate SV call sets from the 10xG simulation, and DELLY and LUMPY to call variants using the WGS simulation. We limited our comparison to only large SVs (>80Kbp for inversions, >40Kbp for duplications) and we required >50% reciprocal overlap between the simulation and the prediction for inversions and the duplicated segments using BEDtools [30]. We also require the inferred insertion breakpoint is within a distance of *µ*_*molecule*_*/*2 (in simulation experiments *µ*_*molecule*_ = 40 Kbp) of the simulation breakpoint to consider a duplication to be correctly predicted.

We present the prediction performance of the tools we tested in Table 1. We found that VALOR_2_ is able to correctly predict >82% of large duplications (inverted and direct combined), and 78% of large inversions, while maintaining 92 *-* 96% precision for duplications and 82% precision for inversions. Long Ranger, the other algorithm that used Linked-Reads, correctly predicted 72% of the inversions with 71% precision. Of the WGS-based tools, DELLY achieved high sensitivity for inversions and it was able to correctly predict 84% of large inversions, however it suffered from very low precision (15%). On the contrary, LUMPY achieved high precision (90%), but it was able to discover only 47% of the simulated inversions. This is likely because neither DELLY nor LUMPY were optimized to find such large inversion events. Overall, VALOR_2_ performed the best in terms of precision and recall balance in the simulation experiment.

**Table 1:** Prediction performance evaluation using simulated structural variants.

| Variant | Tool | # Simulated | # Predicted | True | False | Precision | Recall |
| --- | --- | --- | --- | --- | --- | --- | --- |
| Duplication (direct) | VALOR<sub>2</sub> | 78 | 66 | 61 | 5 | 0.92 | 0.78 |
| Duplication (inverted) | VALOR<sub>2</sub> | 56 | 51 | 49 | 2 | 0.96 | 0.88 |
| Inversion | VALOR<sub>2</sub> | 94 | 65 | 64 | 1 | 0.98 | 0.76 |
|  | LUMPY | 94 | 42 | 44 | 4 | 0.90 | 0.47 |
|  | DELLY | 94 | 896 | 79 | 761 | 0.15 | 0.84 |
|  | Long Ranger | 94 | 92 | 68 | 27 | 0.71 | 0.72 |

We evaluate prediction performance of only large ( $>80$ Kbp for inversions, $>40$ Kbp for duplications) SVs. Note that LUMPY, DELLY, and Long Ranger are not able to call interspersed duplications, thus we provide only the inversion prediction benchmark. Precision is calculated as $\frac{TP}{TP+FP}$ , and recall is defined as $\frac{TP}{TP+FN}$ , where TP: true positive, FP: false positive, FN: false negative.

Here we only focus on large (> 80Kbp) inversions in the NA12878 genome. InvFEST-Valid.: validated inversions in the genome of NA12878, InvFEST-Pred.: predicted inversions in the genome of NA12878, InvFEST-All: all inversions reported in the InvFEST database [25], except those that are annotated as *unreliable prediction*.

### 3.2 NA12878 genome

We also compared the performance of VALOR_2_ with that of Long Ranger on the NA12878 germline genome, along with other commonly used SV callers (DELLY AND LUMPY). NA12878 phased variant calls were obtained from 10X Genomics on their Chromium platform. From these we extracted 476 large inversions, 14 of which were also present in the InvFEST database (Table 2) but only one was experimentally validated. When given the same data, VALOR_2_ was able to call 135 inversions, a higher percentage of which were found in the InvFEST database that also included six experimentally validated inversions. Of the four tools we tested, VALOR_2_ had the largest number of validated inversions within its call set while predicting the second lowest number of total inversions (only LUMPY, which only called 7 inversions, has fewer). This result further highlights the IH: superior precision and recall of VALOR_2_. DELLY was able to identify 24 inversions in the NA12878 genome which were also in the InvFEST database but called a total of 2,340 inversions. A majority of these calls were only predicted by DELLY and due to a lack of precision, may signify an over-representation of false positives (Figure 4). VALOR_2_ was very useful in identifying large scale duplications by exploiting linked read information in the NA12878 sequencing data. We predicted multiple direct segmental duplications and inverted duplications with chromosomes 1 and 16 containing both classes of duplications (Table 3).

**Table 2:** Inversion prediction performance evaluation in the NA12878 genome using InvFEST database.

|  | Called | InvFEST-Valid. | InvFEST-Pred. | InvFEST-All |
| --- | --- | --- | --- | --- |
| VALOR<sub>2</sub> | 135 | 6 | 5 | 17 |
| Long Ranger | 476 | 1 | 10 | 14 |
| LUMPY | 7 | 0 | 0 | 0 |
| DELLY | 2,340 | 1 | 6 | 24 |

**Table 3:** Segmental duplications predicted in the NA12878 genome using VALOR_2_.

| Chr | Start | End | Type | Target | No. of genes |
| --- | --- | --- | --- | --- | --- |
| 1 | 120,600,786 | 120,692,870 | Direct | 1q21.1 | 1 |
| 1 | 144,832,884 | 145,751,706 | Direct | 1p22.3 | 25 |
| 1 | 145,062,336 | 145,116,024 | Direct | 1p11.2 |  |
| 16 | 86,451,165 | 86,498,200 | Direct | 16q11.2 |  |
| 17 | 21,522,544 | 21,551,840 | Direct | 17p11.2 |  |
| 1 | 17,019,657 | 17,111,181 | Inverted | 1q42.3 | 4 |
| 1 | 145,983,326 | 146,027,347 | Inverted | 1p22.3 | 3 |
| 4 | 15,160 | 67,199 | Inverted | 4q35.2 | 2 |
| 8 | 2,189,297 | 2,290,508 | Inverted | 8p23.2 |  |
| 10 | 46,965,140 | 47,022,150 | Inverted | 10q11.22 | 2 |
| 11 | 4,250,956 | 4,331,367 | Inverted | 11p15.4 |  |
| 16 | 21,542,145 | 21,593,639 | Inverted | 16p12.2 |  |
| 16 | 22,543,245 | 22,709,969 | Inverted | 16p12.2 | 2 |
| X | 153,423,995 | 153,485,001 | Inverted | Xq28 | 3 |

**Figure 4:**
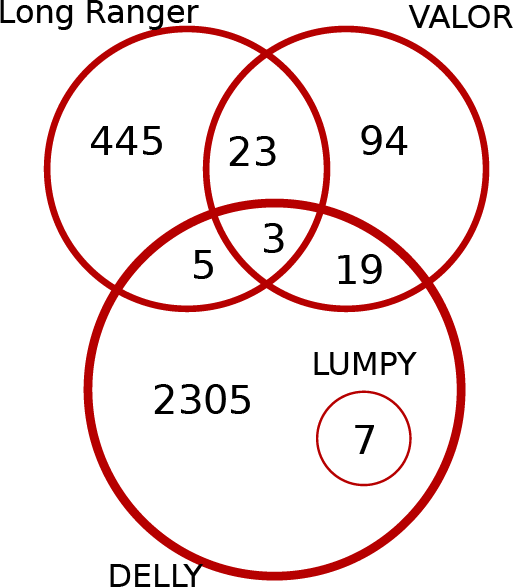
Comparison of the inversion predictions (> 80 Kb) by VALOR_2_, Long Ranger, DELLY, and LUMPY in the NA12878 genome.

### 3.3 CHM1 genome

Finally we tested VALOR_2_ using Linked-Read data set of a haploid human genome cell line (CHM1 [15, 39, 7]). We used VALOR_2_ to find inversions and segmental duplications. Overall, VALOR_2_ characterized 133 inversions (>80 Kbp), 14 inverted and 22 direct segmental duplications (>40 Kb). Unfortunately there are no gold standard data sets for SDs for this genome available in the literature, and the largest previously reported inversion in [7] is 36 Kbp, which is less than the smallest inversion that VALOR_2_ predicts. We therefore compared only with the large inversions in the InvFEST database, and we found that 10% (16/117) of VALOR_2_ predictions were present in InvFEST (Figure 5).

**Figure 5:**
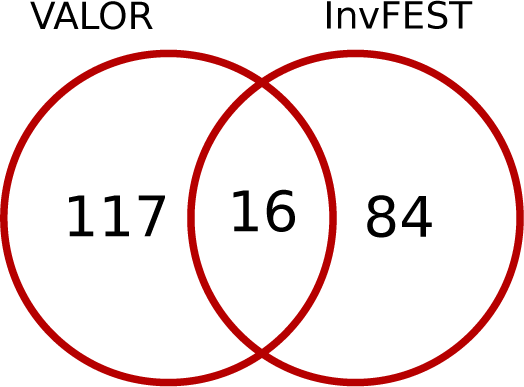
Intersection of all inversions reported by InvFEST (validated or predicted) with VALOR_2_ predictions on CHM1 genome.

## 4 Discussion

In this work, we presented novel algorithms to effectively utilize the encoded long-range information in Linked-Read data for the purpose of characterizing large-scale structural variations. The current state of the art SV detection techniques using Linked-Read such as Long Ranger or GROC-SV are optimized for certain range of SV sizes. For example, GROC-SVs achieves the best sensitivity for events in the range of (30 Kb-100 Kb). However, our technique, VALOR can detect events of a size larger than 100 Kb, including segmental duplications. A future direction for our study is to integrate additional techniques such as local assembly to characterize smaller-scale SVs (i.e. starting from only 50 bp) and to resolve SV breakpoints more precisely. Although single molecule techniques such as Oxford Nanopore (ONT) promise to generate reads in the order of 100 Kb in length, the current error rate, the cost, and the low throughput make them prohibitive in practice. As these techniques get developed and more cost effective, we can explore them not only for the purpose of further validation of our method but also for devising integrative techniques to fully resolve the complexity of repetitive DNA common in mammalian genomes. Another future direction to get more confirmation of our call sets will be using techniques such as fluorescent in situ hybridization (FISH).

## Acknowledgements

We thank H. İ. Özercan, A. Soylev and D. Meleshko for computational support.

## Funding

This work was supported by a grant by TÜBİTAK (215E172), an EMBO Installation Grant (IG-2521), and a Marie Curie Career Integration Grant (303772) to C.A. This work was also supported by start-up funds (Weill Cornell Medicine) to I.H. C.R. received support from the Tri-Institutional Training Program in Computational Biology and Medicine (via NIH training grant 1T32GM083937). The authors also acknowledge the Computational Genomics Summer Institute funded by NIH grant GM112625 that fostered international collaboration among the groups involved in this project.

## Conflict of Interest

None to declare.

## Data availability

The NA12878 genome sequenced with the 10x Genomics Platform is available via the Genome in a Bottle Project [49] FTP site at ftp://ftp-trace.ncbi.nlm.nih.gov/giab/ftp/data/NA12878/10Xgenomics_ChromiumGenome_LongRanger2.1_09302016/NA12878_hg19/ Short read sequencing data for the same genome can be downloaded from the Illumina Platinum Genomes Project at https://www.illumina.com/platinumgenomes.html. The CHM1 genome generated with 10xG Linked-Reads is available at https://support.10xgenomics.com/de-novo-assembly/datasets/2.0.0/chm.

## Footnotes

1 e.g., 30X sequence coverage corresponds to 150X physical coverage.

2 Corresponds to 2 *· µ*_molecule_ and *µ*_molecule_*/*2, respectively.

